# *Af*Rip3, a RIP3-like kinase, is identified as a key modulator of necroptotic death in *Aspergillus fumigatus*

**DOI:** 10.1101/2020.03.25.007617

**Authors:** Jianbo Dai, Linlu Gao, Prakriti Sharma Ghimire, Hui Zhou, Yang Lü, Jinghua Yang, Haomiao Ouyang, Cheng Jin

**Affiliations:** State Key Laboratory of Mycology, Institute of Microbiology, Chinese Academy of Sciences, Beijing 100101, China; University of Chinese Academy of Sciences, Beijing, China

## Abstract

*Aspergillus fumigatus* exhibits autophagic and necroptotic process when its GPI anchor synthesis is suppressed. A putative kinase (AFUA_6G02590) is found to be overexpressed in response to GPI anchor suppression and identified as a RIP3-like protein, namely *Af*Rip3. To elucidate its function, in this study a Af*rip3*-overexpressing strain OE-Af*rip3* was constructed. Although OE-Af*rip3* strain exhibited an increased cell death, neither apoptotic nor autophagic process was activated. Our evidences demonstrated that overexpression of Af*rip3* gene in *A. fumigatus* only led to necroptosis, while the Af*rip3*-knockout mutant was unable to activate necroptotic process. Further analysis revealed that both JNK and SMase pathways were activated in OE-Af*rip3* strain, by which an increase of reactive oxygen species (ROS) was induced. We also showed that expression of Af*rip3* gene was induced by Ca^2+^. In addition, eEF1Bγ and adenylylsulfate kinase (ASK) were identified as potential candidates to interact with *Af*Rip3. These results indicate that *Af*Rip3 is a key modulator that activates necroptotic process in *A. fumigatus*, which can be induced by Ca^2+^ and in turn activate JNK (c-Jun NH_2_-terminal kinase) and SMase (sphingomyelinase) pathway. Our findings suggest that necroptotic pathway in *A. fumigatus* is distinct from that in mammalian cell and may provide a new strategy for development of anti-fungal drug.

**Author summary:** *Aspergillus fumigatus* is a human fungal pathogen and causes invasive aspergillosis (IA) in immunocompromised patients with high mortality (30-95%). Development of novel therapies is urgently needed. In this study, we confirm *Af*Rip3 (AFUA_6G02590), a RIP3-like protein, is a key modulator that activates necroptotic process in *A. fumigatus*. We also find that cytosolic Ca^2+^ can induce the expression of Af*rip3* and activated *Af*Rip3 in turn activate JNK (c-Jun NH_2_-terminal kinase) and SMase (sphingomyelinase) pathway. Our findings suggest that necroptotic pathway in *A. fumigatus* is distinct from that in mammalian cell and may provide a new strategy for development of anti-fungal drug.

## Introduction

Programmed cell death (PCD) plays a significant role in the development, immune homeostasis, and host defense of multicellular organisms. To date, three types of PCD have been described in higher eukaryotes, including apoptosis, autophagy, and necroptosis (also known as programmed necrosis). Apoptosis is the most conserved form of PCD, requires the activation of caspases, and is defined by chromatin condensation, DNA fragmentation, cell shrinkage, blebbing of plasma membrane and formation of apoptotic bodies. Autophagy is a lysosome degradation pathway by which cells capture intracellular proteins, lipids and organelles, and deliver them to the lysosome compartment. It is induced under conditions of nutrient starvation, liberating energy stores and promoting cellular survival [1]. Necroptosis, which is previously known as necrosis and once thought to be genetically uncontrolled, is also programmed cell death [2-5] and morphologically characterized by rupture of plasma membrane and organelle breakdown [6-7].

Necroptosis can be initiated by death ligands, Toll-like receptor ligands (TLRs), or microbial infection [8]. The most well-studied signaling pathway induced by death ligands is tumor necrosis factor (TNF). Signaling from TNF receptors activates the receptor-interacting protein kinase 1 (RIP1) and RIP3. RIP1 is a death-domain-containing kinase containing a conserved kinase domain in the N-terminus and a RIP homotypic interaction motif (RHIM) domain in the C-terminus, but its kinase activity is dispensable for inducing death-receptor-mediated apoptosis [9-10]. RIP3 shares 30%-40% sequence similarity with RIP1 and is essential for necroptosis. RIP1, RIP3 and mixed lineage kinase domain-like protein (MLKL) form a necrosis signaling complex named necrosome, within which MLKL is phosphorylated by RIP3. The phosphorylated MLKL forms an oligomer and binds to the plasma and intracellular membranes to form membrane-disrupting pores, which results in necroptotic death [10-14]. On the other hand, RIP3-dependent necrosis can also proceed without RIP1. Indeed, RIP3-dependent necroptosis upon ectopic expression of RIP3 has been described in RIP1-deficient MEF cells [15]. Under certain cellular conditions necroptosis can occur in the absence of RIP1 in L929 cells [16]. However, the precise mechanism of RIP1-independent necroptosis remains unclear.

*Aspergillus fumigatus* is a human fungal pathogen capable of causing infections ranging from allergic to invasive disease [17], and the major cause of invasive aspergillosis (IA) in immunocompromised patients [18]. In these patients, the crude mortality is 30-95%. Despite some effective drug treatments, mortality from fungal infections remains about 50%, and new drugs are urgently needed due to the inefficacy, side effects and resistance that have emerged as important factors limiting successful clinical outcome [19-23]. A major barrier for the development of novel therapies is the general lack of capacity in fungal pathogen research [24].

Although PCD has already been discovered from bacteria to animals [25-28], in contrast to that in mammalian cells, little is known about death process in filamentous fungi. In filamentous fungi autolysis is a highly regulated and natural process that occurs later in older, stationary phase cultures and leads to the progressive disintegration of the mycelium. In *A. fumigatus*, it has been revealed that autolysis during the stationary phase is an apoptotic process and caspase-dependent [29]. However, autophagic and necroptotic process in *A. fumigatus* remain unclear.

Previously, we have shown that suppression of GPI anchor synthesis leads to both autophagic and necroptotic process in *A. fumigatus*. Suppression of the GPI anchor synthesis leads to activation of phosphatidylinositol (PtdIns) signaling and ER stress, which in turn induce increased cytosolic Ca^2+^, activate PtdIns3K and induce autophagy [30]. Meanwhile an necroptotic process is also induced. Although the mechanism of necroptosis remains unclear, a putative kinase (AFUA_6G02590) has been identified as a RIP3-like protein, namely *Af*Rip3, which only contains a kinase domain and lacks the homotypic interaction motif (RHIM) required for interaction with RIP1 [30].

To elucidate the potential role of *Af*Rip3 in cell death process of *A. fumigatus*, a Af*rip3-*overexpressing strain OE-Af*rip3* and a Af*rip3*-knockout mutant were obtained in this study. Analysis of the OE-Af*rip3* strain revealed that overexpression of the Af*rip3* only induced necroptosis but not apoptosis or autophagy. Meanwhile the Af*rip3-*knockout strain was unable to activate necroptotic process. Further analysis revealed that *Af*Rip3 was activated by calcium and executed necroptosis by activating JNK and SMase pathway. In addition, eEF1Bγ and adenylylsulfate kinase (ASK) were identified as potential candidates to interact with *Af*Rip3.

## Results

### Af*rip3*-overexpression induces cell death of *A. fumigatus*

Expression vector was constructed by introducing a copy of Af*rip3* gene into pVG2.2, a vector comprising of two modules: one module ensures constitutive expression of the tetracycline dependent transactivator rtTA2S-M2 and another one harbors the rtTA2S-M2-dependent promoter that controls expression of the gene of interest [31-32]. The Af*rip3* expression vector was then transformed into *A. fumigatus* and screened for uridine and uracil autotrophy [33]. As a result, twenty-four transformants were obtained, while sixteen were confirmed to be correct by PCR analysis. As shown in Fig 1A, a 863-bp fragment of *pyrG* and a 1102-bp fragment of pVG2.2 vector were amplified from the genomic DNA of OE-Af*rip3* strain and sequenced, while no such fragments were amplified from the wild-type (WT). Quantitative RT-PCR analysis revealed that the expression of Af*rip3* in OE-Af*rip3* strain was 3.8 times of that in the WT, indicating that OE-Af*rip3* strain was successfully constructed (Fig 1B).

**Fig 1.**
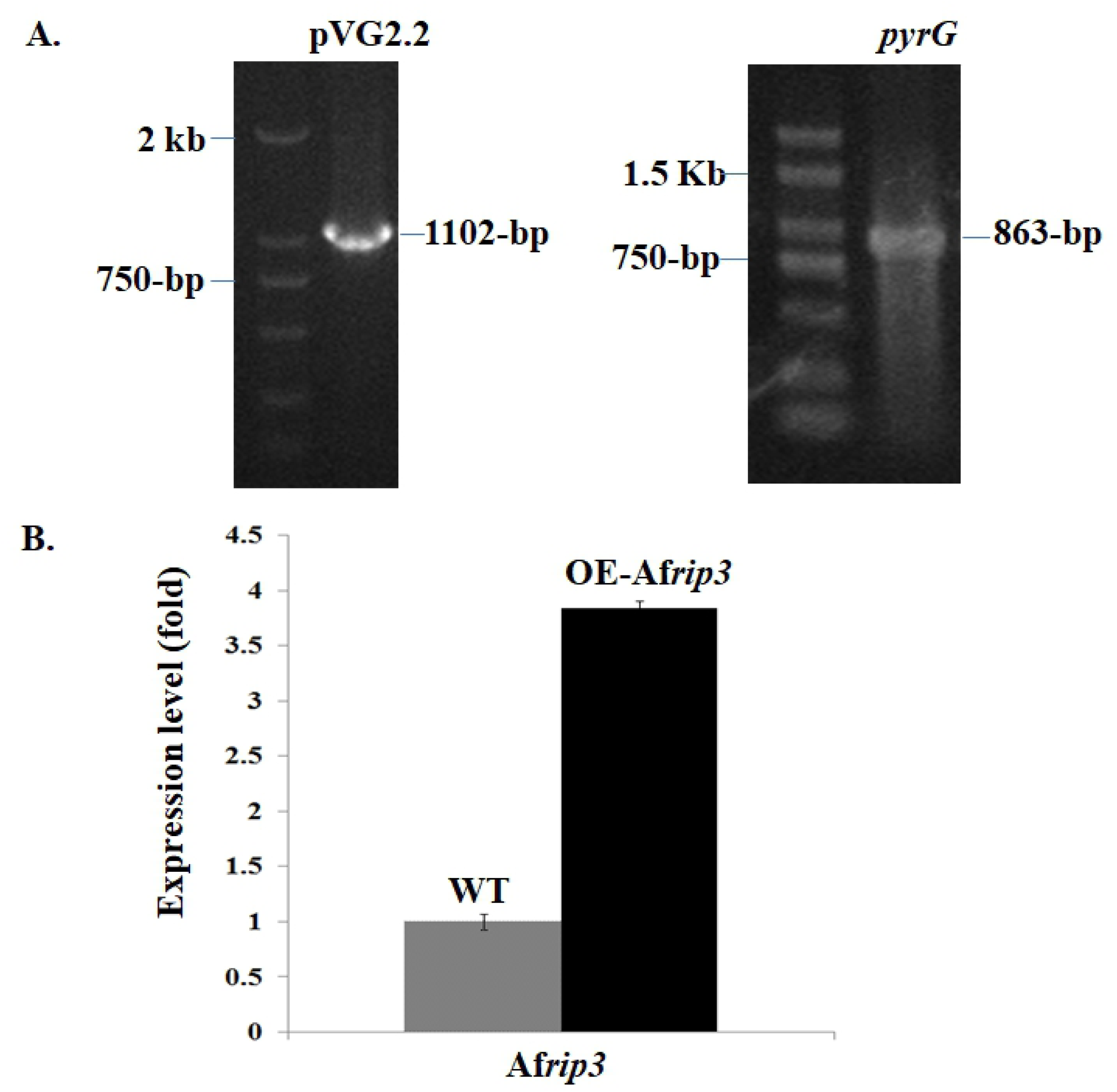
Construction of OE-Af*rip3* strain. In A, PCR confirmation of the OE-Af*rip3* strain was carried by using a primer pair of 5’-ATAGGGCATATTCAACTACCTGGCT-3’ and 5’-GTTTATAGACTCTCAATTCGCGATC-3’ to amplify an 863-bp fragment of the Af*rip3* gene as described under Materials and Methods; in B, quantitative RT-PCR was carried out by using 1 μg RNA isolated from strains as described under Materials and Methods. Quantification of mRNA was performed using the 2^−ΔΔct^ method. Triplicates of samples were analyzed in each assay, and each experiment was repeated at least three times. Results are presented as mean ± SD.

After 24 h of incubation, both WT and OE-Af*rip3* strain reached their log-phase and mycelia were stained with propidium iodide (PI), a dye that labels the nucleus in dying cells and is widely used to detect cell membrane integrity and cell viability [34-36]. As shown in Fig 2, under microscope about 75.5% of mycelia of the the OE-Af*rip3* were PI-positive, while PI staining was not observed in the WT, indicating an occurrence of cell death in OE-Af*rip3* strain at its log-phase. This result indicates that elevated expression of the Af*rip3* induces cell death of *A. fumigatus*.

**Fig 2.**
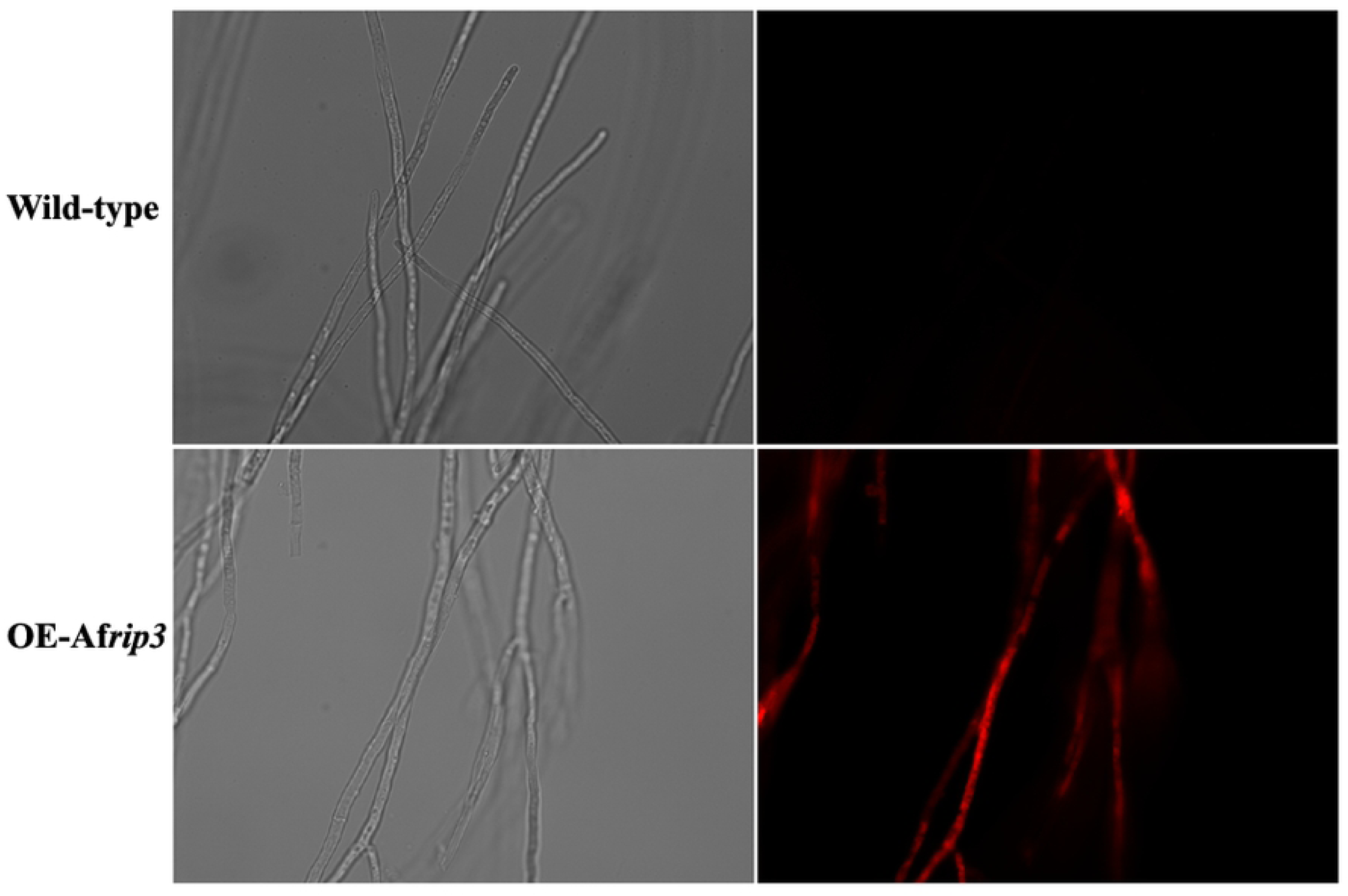
Detection of cell death in OE-Af*rip3* strain. Strains were cultured at 37°C for 24 h. The mycelia were collected by centrifugation at 8,000 rpm at 4°C. The supernatant was thoroughly removed. Sterile filtered solution containing 30 µg/mL propidium iodide (PI) in phosphate-buffered saline (137 mM NaCl, 2.7 mM KCl, 4.3 mM Na_2_HPO_4_•7H_2_O, and 1.4 mM KH_2_PO_4_) was added to the tube. After standing at room temperature for 5 min, the hyphae were washed by PBS buffer for 3 times and visualized under the fluorescent microscope.

### Analysis of PCD pathway in strain OE-Af*rip3*

To clarify the pathway of cell death in OE-Af*rip3* strain, three types of PCDs were analyzed in this study. In mammalian cells apoptosis is featured with activation of Caspase-8 and translocation of phosphatidylserine (PtdSer) [37-38]. As apoptosis in *A. fumigatus* shares features of the apoptotic pathway of mammalian cells [29], we first checked CasA, a counterpart of caspase-8, and exposure of PtdSer in OE-Af*rip3* strain. When the WT was cultured in presence of apoptosis-inducer dexamethasone, an elevated activity of CasA was induced (Fig 3A) and exposure of PtdSer was detected (Fig 3B), indicating an occurrence of dexamethasone-induced apoptosis in *A. fumigatus*. However, as compared with that in WT, the CasA activity in OE-Af*rip3* strain was declined by 39.3% (Fig 3A) and exposure of PtdSer was not detected (Fig 3B). These results demonstrate that overexpression of the Af*rip3* does not activate apoptotic pathway in *A. fumigatus*.

Atg8/LC3 (microtubule-associated protein 1 light chain 3) is a reliable markers for autophagy [39]. Previously, we have shown that suppression of GPI anchor synthesis induces activation of Atg8/LC3 homolog and autophagy [30]. To determine if the autophagic pathway was activated in OE-Af*rip3* strain, Atg8/LC3 homolog was detected by either anti-LC3I and anti-LC3II antibodies. As shown in Fig 4, both LC3I and LC3II detected in OE-Af*rip3* strain were similar with that in the WT. This result demonstrates that overexpression of the Af*rip3* gene does not elicit autophagic process.

**Fig 3.**
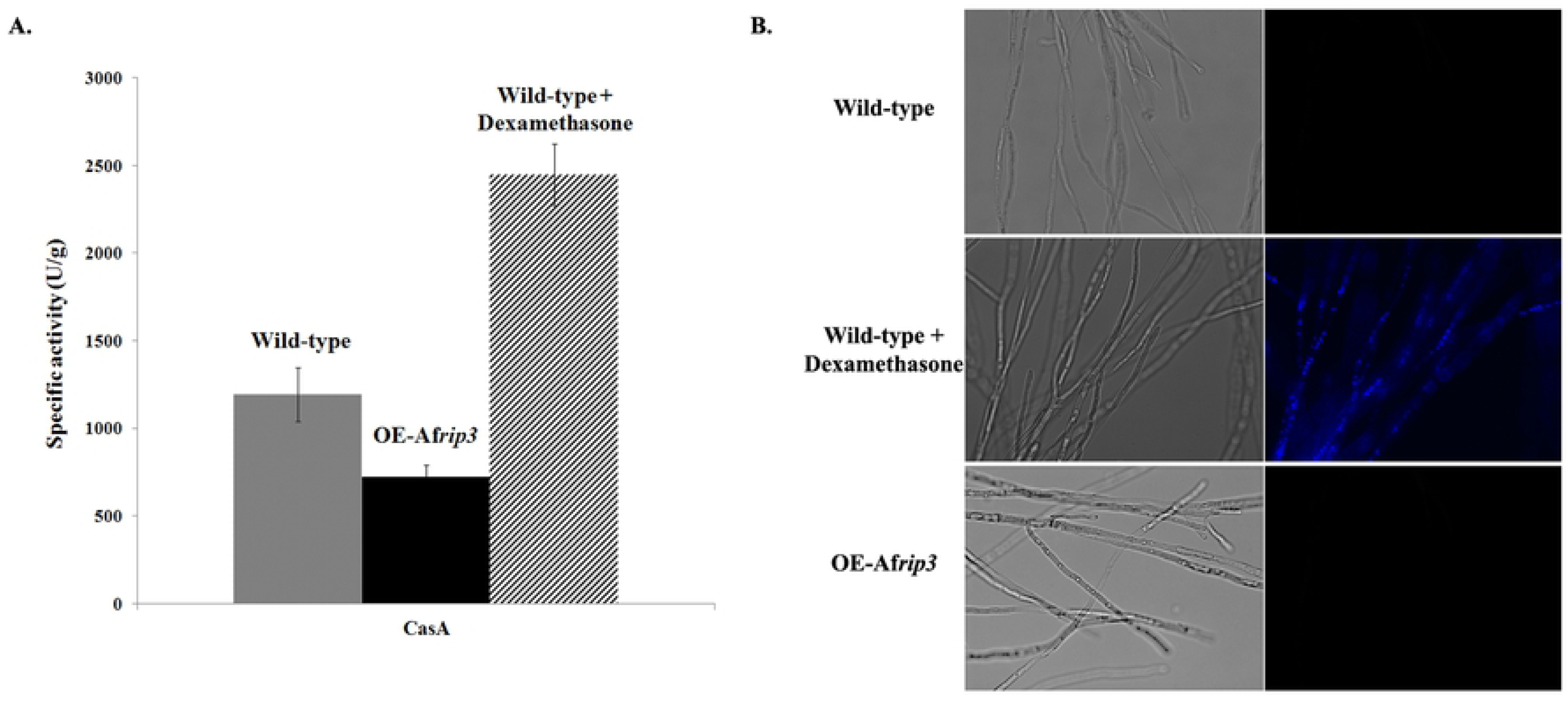
Determination of CasA activity and surface-exposed phosphatidylserine. In A, strains were cultivated with or without dexamethasone. Intracellular proteins were extracted for activity assay. Caspase activity was determined using Caspase Fluorescent (AMC) Substrate/Inhibitor QuantiPakTM (BioMol International). The results are represented as the mean ± SD of three replicates. In B, after cultured in CM for 24 h, the mycelia were collected and washed with PBS three times. The mycelia were resuspend in 500 µL PBS and detected with Annexin V-FITC Apoptosis Detection Kit (Sigma) under microscope (Zeiss).

**Fig 4.**
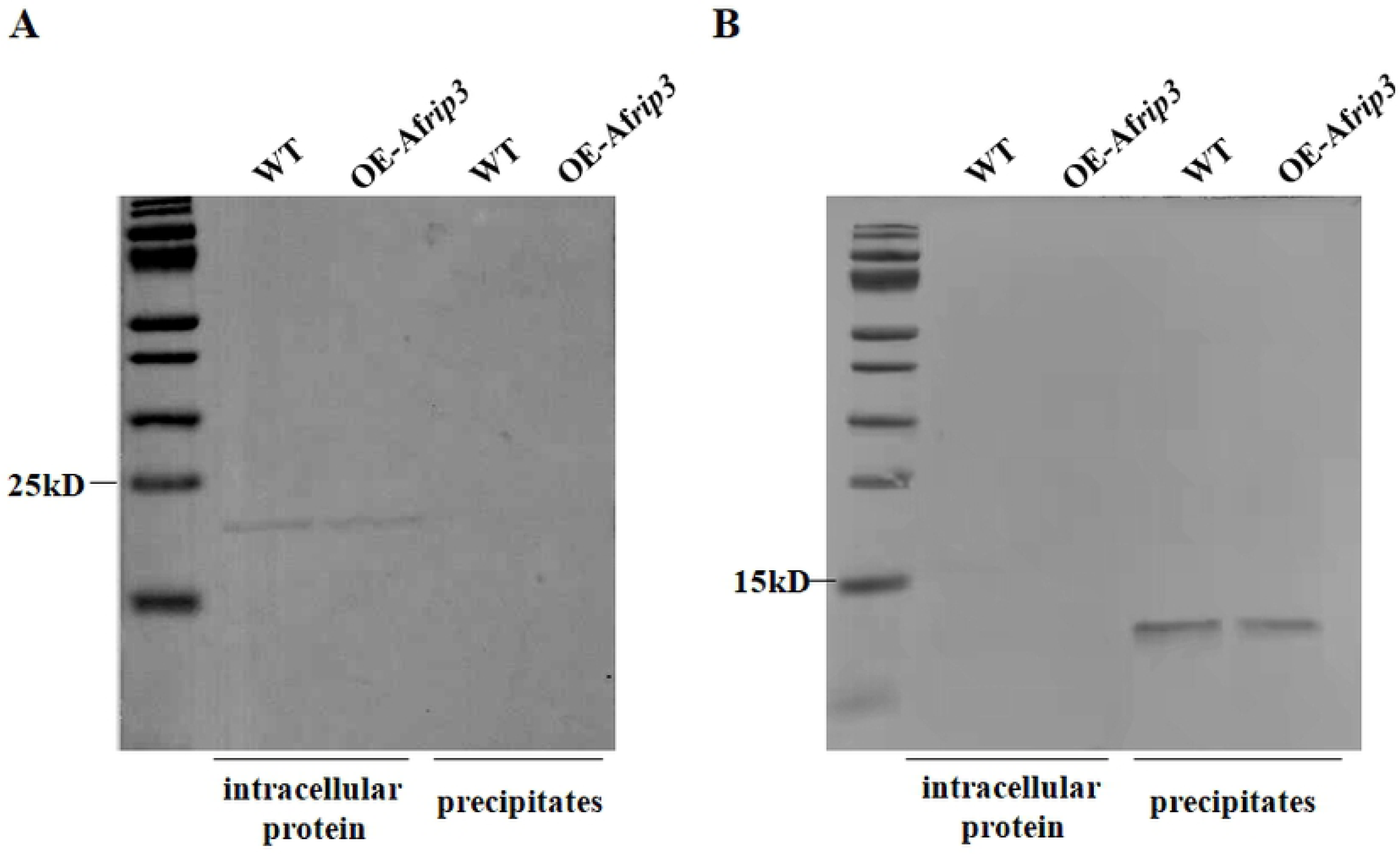
Determination of LC3 in OE-Af*rip3* strain. Proteins were separated on SDS-PAGE and transferred onto PVDF membranes (Millipore). The membrane was blocked with 5% fat-free milk in TBST for 2 h at room temperature, and then incubated with anti-LC3I (A) or anti-LC3II (B) antibody (Sigma) at 4°C overnight. Then the membrane was washed three times with TBST buffer and incubated with an AP-conjugated secondary antibody for 1 h at room temperature. After washing three times with TBST buffer, bands were detected with NBT/BCIP reagent.

As necroptosis is morphologically characterized by organelle damage, cell swelling and rupture of the plasma membrane [8], we further checked the morphology of OE-Af*rip3* strain under transmission electron microscope (TEM). As shown in Fig 5, massive vacuolization and translucent cytoplasma were observed in 85.2% cells of the OE-Af*rip3* grown in CM for 24 h, while no such phenotype was found in the WT. This observation suggests a release of the cytoplastic contents in the OE-Af*rip3* cells. To evaluate the rupture of the plasma membrane of OE-Af*rip3* strain, the leakage of a cytoplasmic lactate dehydrogenase (LDH) was detected by using the method established by Xie et al. [40]. As shown in Fig 6A, LDH activity in the WT was determined as 63 U/mg, while LDH activity in OE-Af*rip3* strain was 152U/mg, which is 2.4-fold of that in the WT. We further examined plasma membrane integrity in OE-Af*rip3* strain by using GPI-anchored membrane protein Ecm33 as a reporter. Western blotting analysis revealed that Ecm33 was increased in the culture supernatant of OE-Af*rip3* strain (Fig 6B). These results confirm an occurrence of membrane rupture and a significant leakage of intracellular enzyme and membrane proteins in OE-Af*rip3* strain, indicating an activation of necroptotic process in OE-Af*rip3* strain.

**Fig 5.**
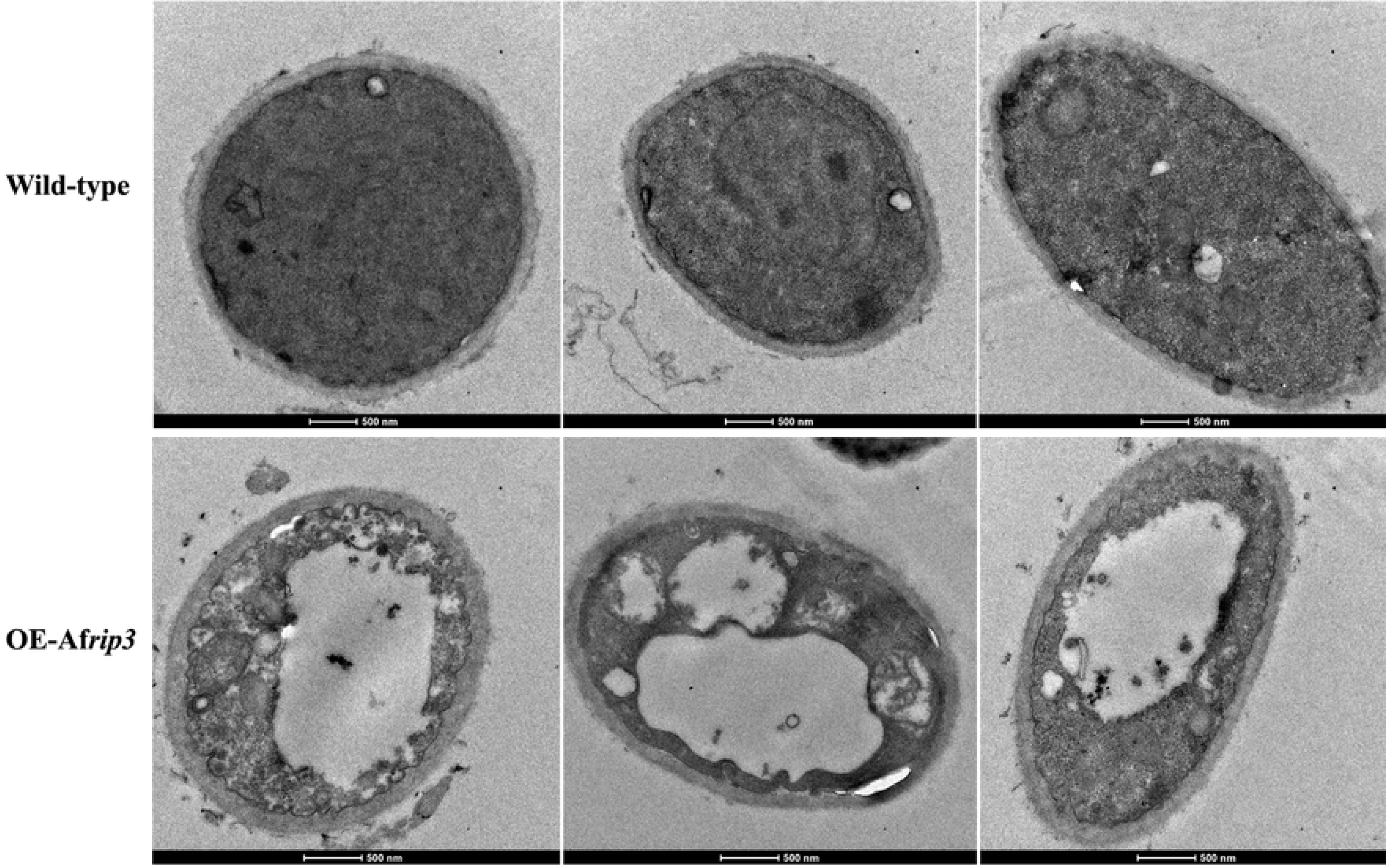
Transmission electron microscopy of OE-Af*rip3* strain. After cultivation in CM for 24 h, mycelia were collected, suspended in PBS and fixed overnight at 4°C in 2.5% (w/v) glutaraldehyde. The samples were post-fixed with 1% (w/v) osmium tetroxide solution for 2h at room temperature, dehydrated in an acetone series (30, 50, 70, 85, 95 and 100%) and subjected to 2% uranyl acetate and 30% methanol. Samples were embedded in Spurr’s plastic and sectioned with a diamond knife. Thin sections were placed on copper grids and stained with uranyl acetate and lead citrate, and examined under an FEI Tecnai Sprit transmission electron microscope (FEI, Hillsboro, OR, USA)

**Fig 6.**
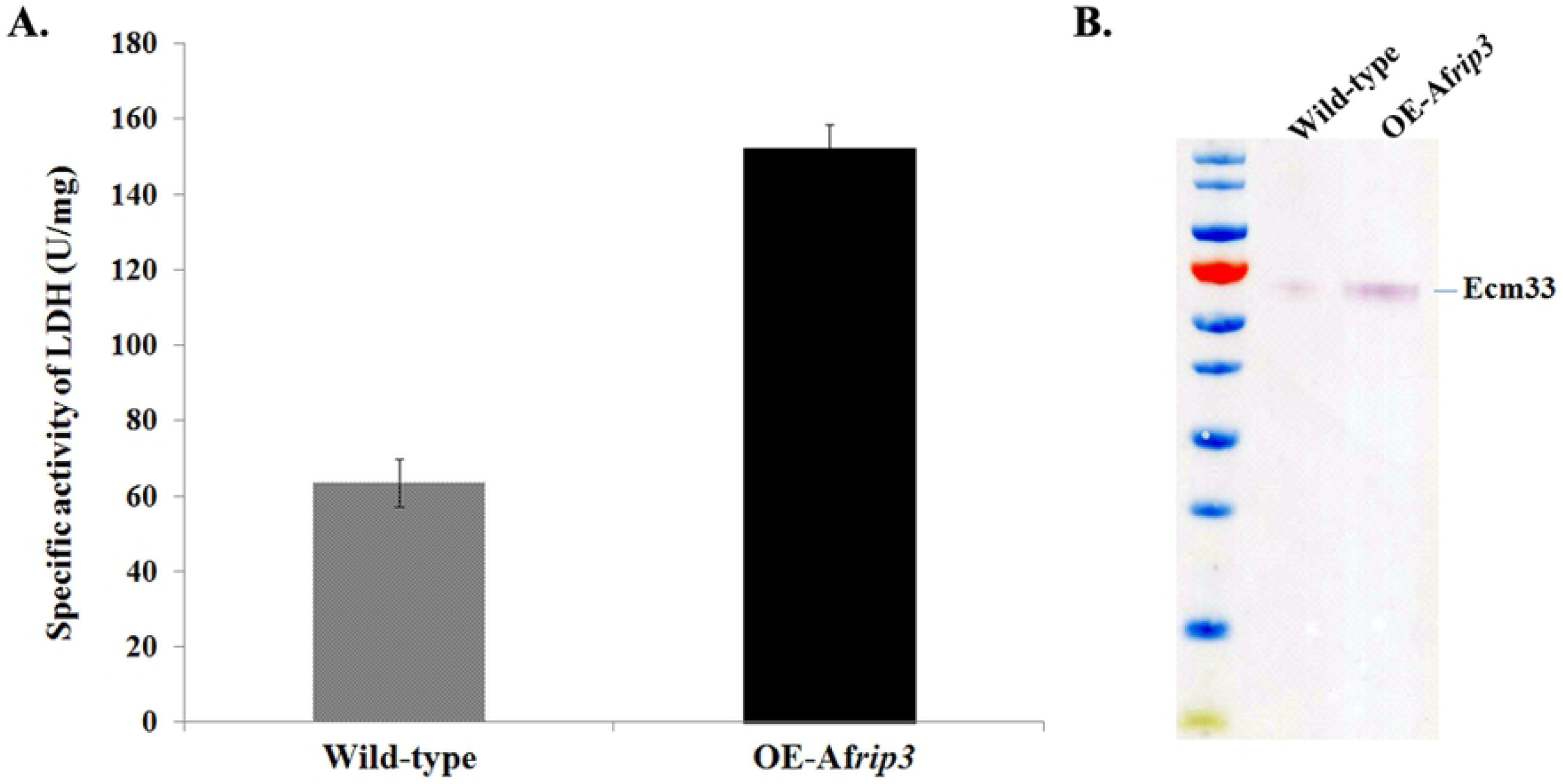
Release of intracellular lactate dehydrogenase and membrane protein Ecm33 in OE-Af*rip3* strain. Extracellular proteins were extracted as described under Material and Methods. In A, the activity of released lactate dehydrogenase (LDH) was determined by LDH Release Assay Kit (Beyotime, China). The absorbance was read at 490 nm. The results are represented as the mean ± SD of three replicates. In B, Extracellular proteins from either WT or OE-Af*rip3* strain were separated on SDS-PAGE and transferred onto PVDF membranes (Millipore). The membrane was blocked with 5% fat-free milk in TBST for 2 h at room temperature, and then incubated with anti-Ecm33 antibody at 4°C overnight. Then the membrane was washed three times with TBST buffer and incubated with an AP-conjugated secondary antibody for 1 h at room temperature. After washing three times with TBST buffer, bands were detected with NBT/BCIP reagent.

To further confirm the regulatory role of *Af*Rip3 in necroptosis, Af*rip3* gene was deleted in *A. fumigatus*. When the WT was cultured in liquid CM supplemented with TSZ, a cocktail of necroptosis-inducers consisting of TNF-α, SM-164 and Z-VAD-FMK, necroptotic cell death was induced. However, in the ΔAf*rip3* mutant no cell death was induced by TSZ (Fig 7). Taken together, our results confirm that *Af*Rip3 is a regulator of necroptotic cell death in *A. fumigatus*.

**Fig 7.**
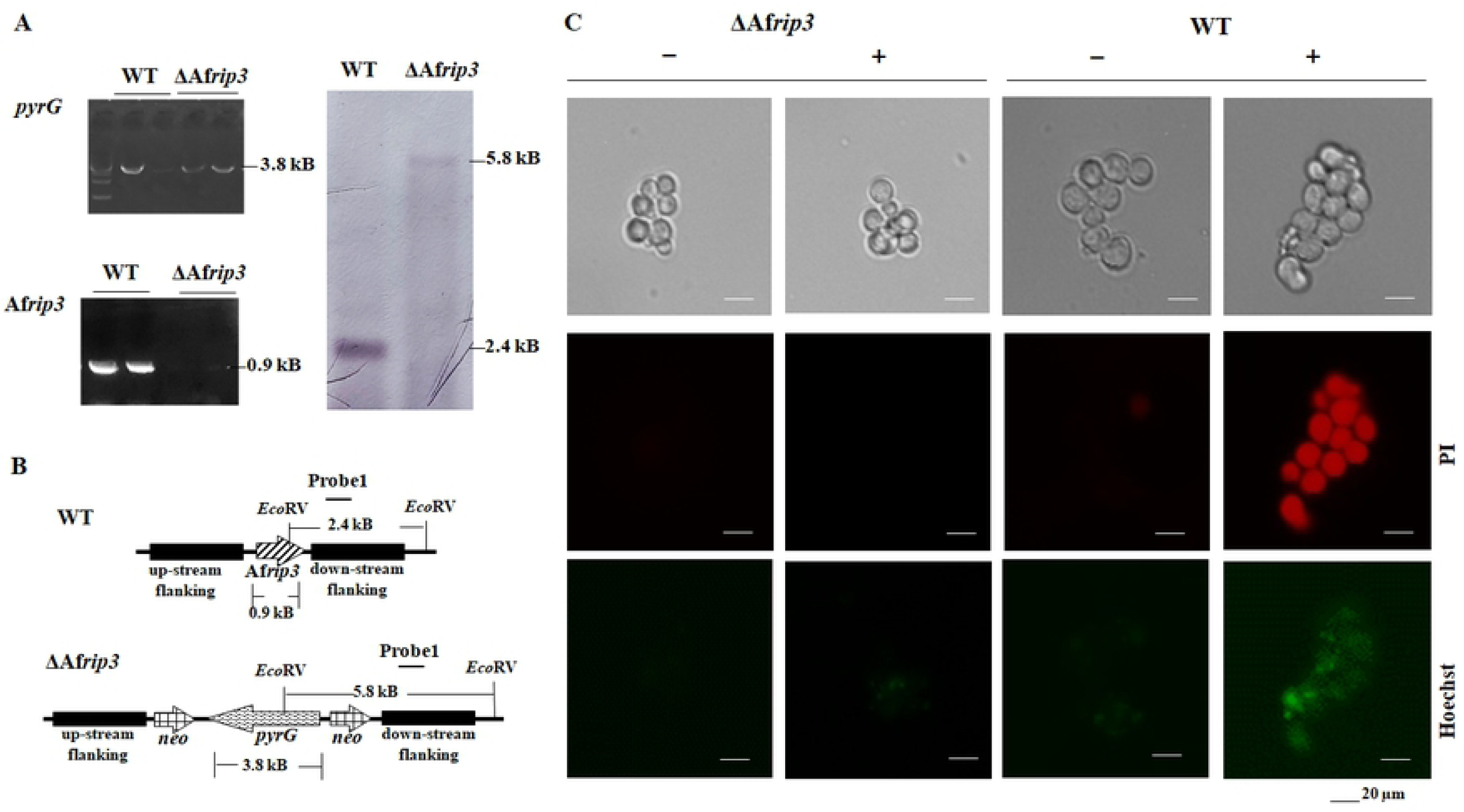
Construction and analysis of the ΔAf*rip3* mutant. The null mutant ΔAf*rip3* was constructed by replacing of Af*rip3* gene with *pyrG* as described under Matherials and Methods. The mutant was confirmed by PCR and Southern blot (A and B). 1×10^6^ of the WT or mutant spores were inoculated into 1 ml liquid CM with (+) or without (-) 1 μl TSZ and incubated at 37°C for 4 hours, stained with 5 μl Hoechst33342 and 5 μl propidium iodide (PI) at 4°C for 20 minutes (Apoptosis and Necrosis Assay Kit, C1056, Beyotime, China), and then examined under the fluorescence microscope (Zeiss Imager A2, Zeiss, Japan). WT: wild-type; PI: propidium iodide; TSZ, mixture of TNF-α, SM-164 and Z-VAD-FMK (Necroptosis Inducer Kit, C1058S, Beyotime, China). Scale bar: 20μm.

### Regulation of Af*rip3* gene expression

As release of the ER-Ca^2+^ is the main effect causes both autophagy and necroptosis in *A. fumigatus* [30], we assumed that Ca^2+^ is also an factor to induce expression of the Af*rip3*. To verify this hypothesis, we tested the effect of calcium ion on expression of the Af*rip3*. As shown in Fig 8A, when the WT was cultured with CaCl_2_ for 24 h, the expression of Af*rip3* gene was up-regulated up to 2.5-, 1.25- and 1.5-fold in presence of 0.1 mM, 1 mM and 10 mM CaCl_2_, respectively. We also tested the effect of inositol and fermentation broth on expression of the Af*rip3*. Inositol was able to slightly up-regulate the expression of the Af*rip3*, while fermentation broth slightly inhibited the expression of the Af*rip3*. These results demonstrate that the expression of Af*rip3* gene is activated by Ca^2+^, but not by inositol or metabolites produced by aging *A. fumigatus*.

**Fig 8.**
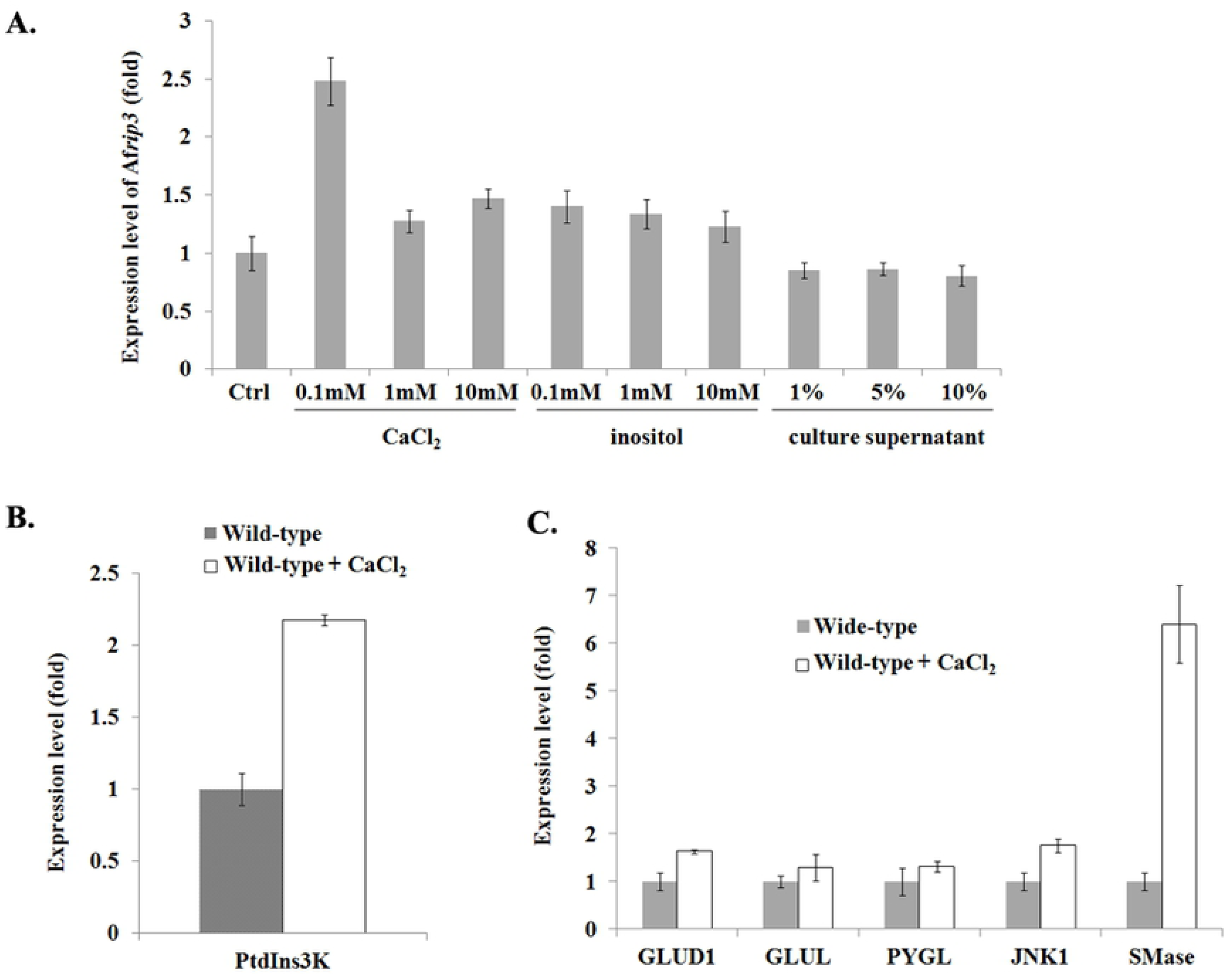
Effect of Ca^2+^ on the programmed cell death (PCD) of *A. fumigatus*. 2×10^8^ spores were cultured in culture medium supplied with CaCl_2_ at 37°C for 24 h. RNAs were extracted and quantified as described under Materials and Methods. In A, the expression levels of Af*rip3* were determined with RNAs extracted from the OE-Af*rip3* strain cultured with different concentrations of CaCl_2_; in B, expression levels of glutamate dehydrogenase1 (GLUD1), glutamate-ammonia ligase (GLUL), glycogen phosphorylase (PYGL), c-Jun NH_2_-terminal kinase 1 (JNK1) and sphingomyelinase (SMase) were determined with RNAs extracted from the OE-Af*rip3* strain cultured with 0.1 mM of CaCl_2_, respectively; and in C, expression level of the gene encoding Vps34/PtdIns3K was determined with RNAs extracted from the OE-Af*rip3* strain cultured with 0.1 mM of CaCl_2_. The results are represented as the mean±SD of three replicates.

During autophagic process, the Vps34-Atg6/beclin1 class III phosphoinositide 3-kinase (PtdIns3K) complex is another important subgroup of the “core” Atg proteins [41]. RT-qPCR analysis showed that Ca^2+^ was able to induce an increased expression of PtdIns3K/Vps34 up to 2.2-fold in the wild-type *A. fumigatus* (Fig 8B). These results are consistent with our previous findings that autophagy in the *afpig-a* conditional mutant is induced by Ca^2+^ [30] and indicate that Ca^2+^ is also able to activate necroptotic process through induction of Af*rip3*-expression in *A. fumigatus*.

### Activation of JNK and SMase pathway by *Af*Rip3

Upon suppression of the GPI anchor synthesis some of the molecules involved in necroptosis are induced at least 1.5-fold, such as glycogen phosphorylase (PYGL), glutamate-ammonia ligase (GLUL), glutamate dehydrogenase 1 (GLUD1), Nfr1/AIF, cyclophilins, JNK1, Hsp70 family proteins, PKA, glyoxalase family proteins and Rab7. While several other proteins required for necroptosis were suppressed, such as sphingomyelin phosphodiesterase (SMase), ceramidase, poly(ADP)-ribose polymerase (PARP) and calpains [15, 30]. In this study, we also tested the expression of these genes in OE-Af*rip3* strain. As summarized in Table 1, Nfr1/AIF, cyclophilins, JNK1, Hsp70 family proteins, glyoxalase family proteins and Rab7 were induced (Table 1), which is consistent with that in the *afpig-a* conditional mutant [30]. On the other hand, although Ca^2+^ was able to induce an elevated expression of the genes encoding PYGL, GLUL, GLUD1, JNK1 and SMase in the wild-type *A. fumigatus* (Fig 8C), PYGL, GLUL, GLUD1, PKA, and Rab7 were suppressed in OE-Af*rip3* strain (Table 1).

**Table 1.**
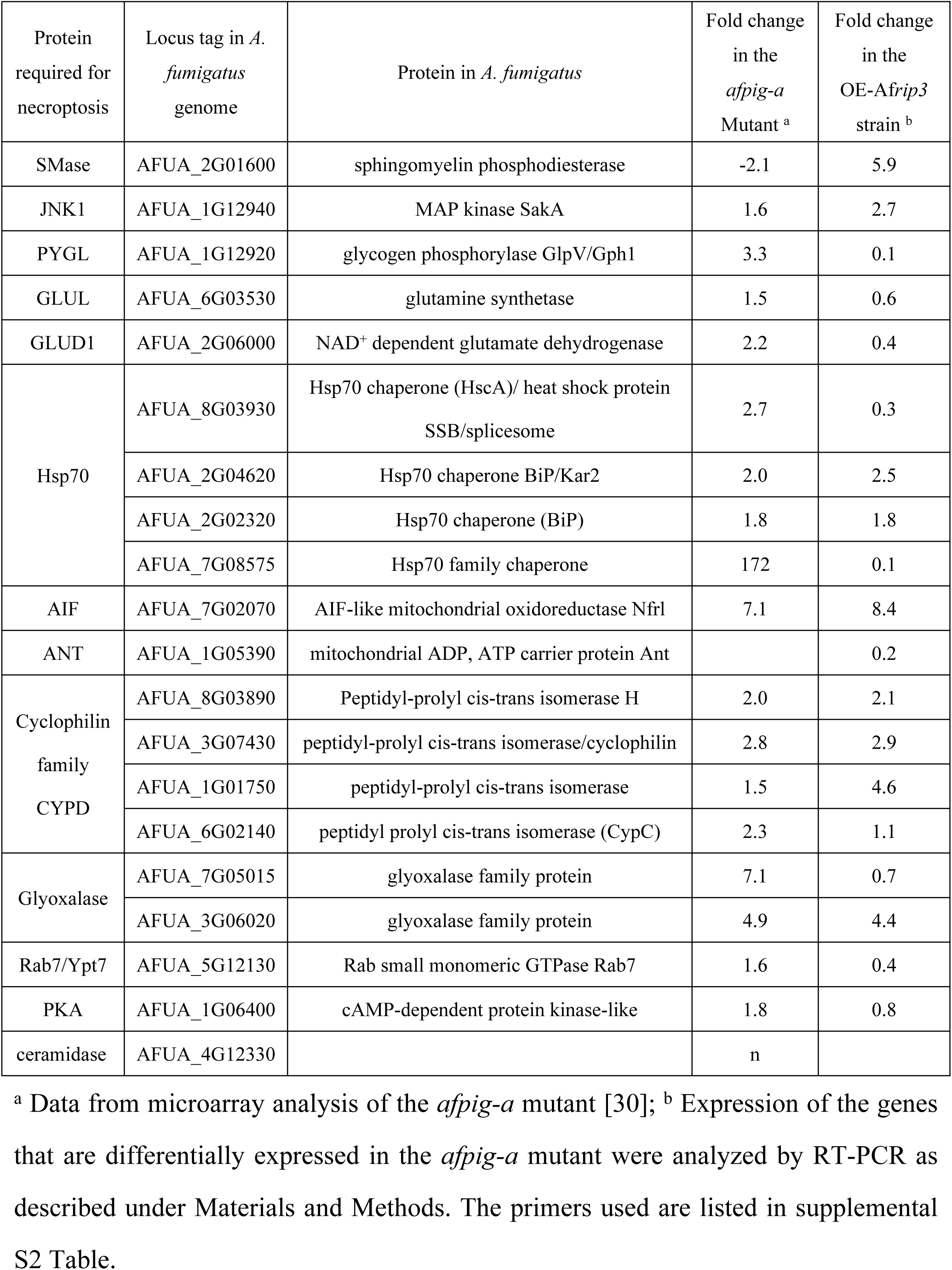
Expression of necroptosis-related genes in OE-Af*rip3* strain.

In mammalian cells, the c-Jun NH_2_-terminal kinase (JNK) is activated by TNF and initiates necroptotic cell death by inducing ROS production [42-44] and sphingomyelinase (SMase) pathway, which lead to lysosomal membrane permeabilization [45-47]. SMase and ceramidase are key enzymes in sphingomyelinase (SMase) pathway. It is interesting to note that, unlike that in the *afpig-a* conditional mutant, SMase and ceramidase in OE-Af*rip3* strain were induced 5.9- and 3.7-fold, respectively (Table 1). As both JNK1 and SMase elicit ROS generation [48], we further detected ROS in the OE-Af*rip3* strain with dihydroethidium, a dye that can permeate viable cells and accumulate in the nucleus when dehydrogenated to ethidium bromide. As shown in Fig 9, OE-Af*rip3* strain was positive-stained by dihydroethidium, whereas the WT was negative, indicating a significant increase of ROS in OE-Af*rip3* strain. All these data establish that overexpression of Af*rip3* gene triggers necroptotic process by activating JNK1 and SMase, thereby allowing the generation of ROS, and further promoting lysosome membrane permeabilization.

**Fig 9.**
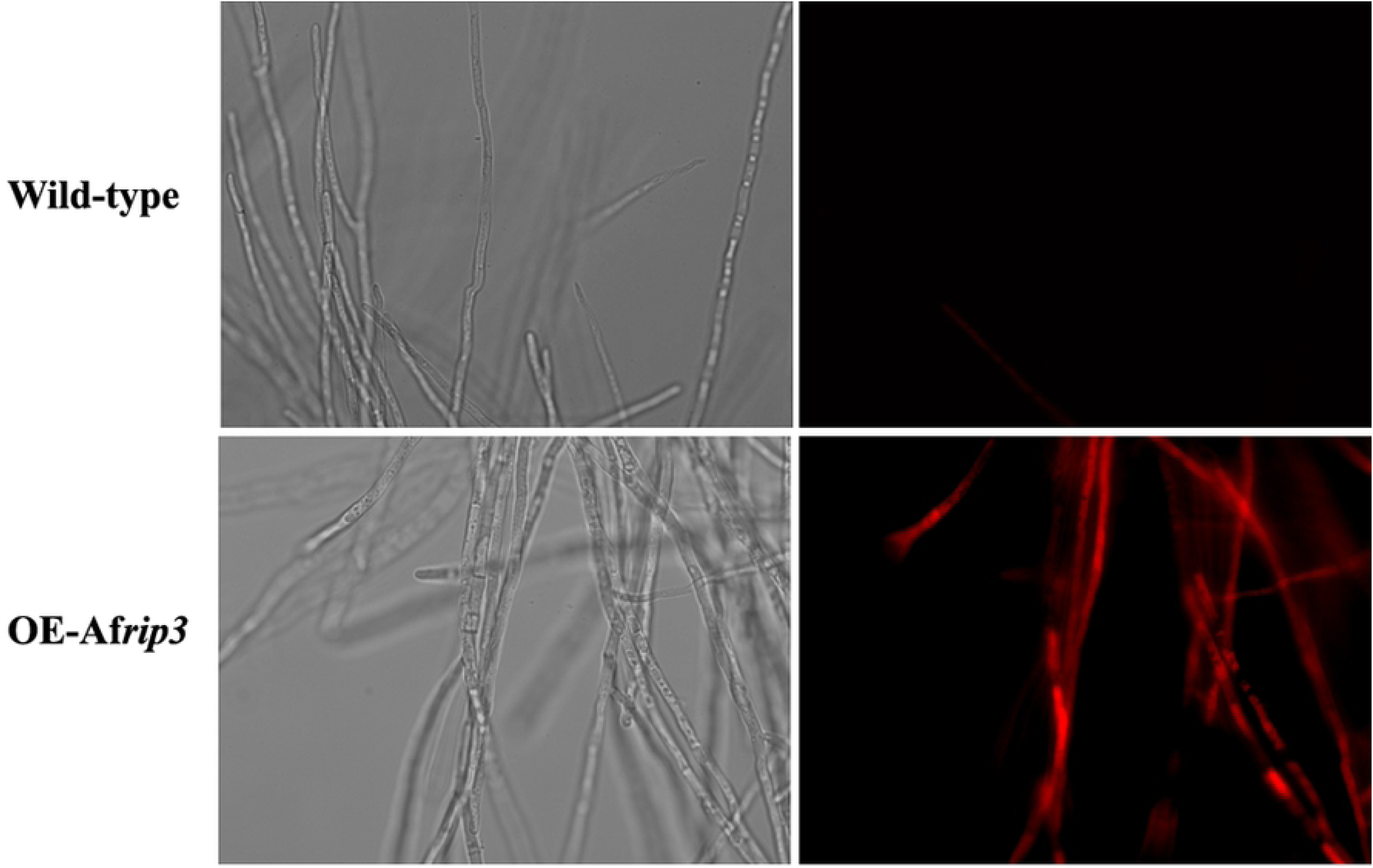
Detection of ROS in OE-Af*rip3* strain. The WT and OE-Af*rip3* strain were cultured in CM for 24h, stained by dihydroethidium and visualized under fluorescence microscope.

### Identification of proteins that potentially interact with *Af*RIP3

In attempt to identify the downstream target of *Af*Rip3, pull-down assay was carried out by using GST-*Af*Rip3 protein expressed in *A. fumigatus* (Fig 10A). A 25 kDa protein band was detected on SDS-PAGE (Fig 10B) and analyzed by mass spectrum. As a result, 23 proteins were identified (S1 Table). Among these proteins, eEF1Bγ and adenylylsulfate kinase (ASK) were confirmed to interact with *Af*Rip3 by co-IP with anti-His-tag mAb-Magnetic Beads (Fig 10C).

**Fig 10.**
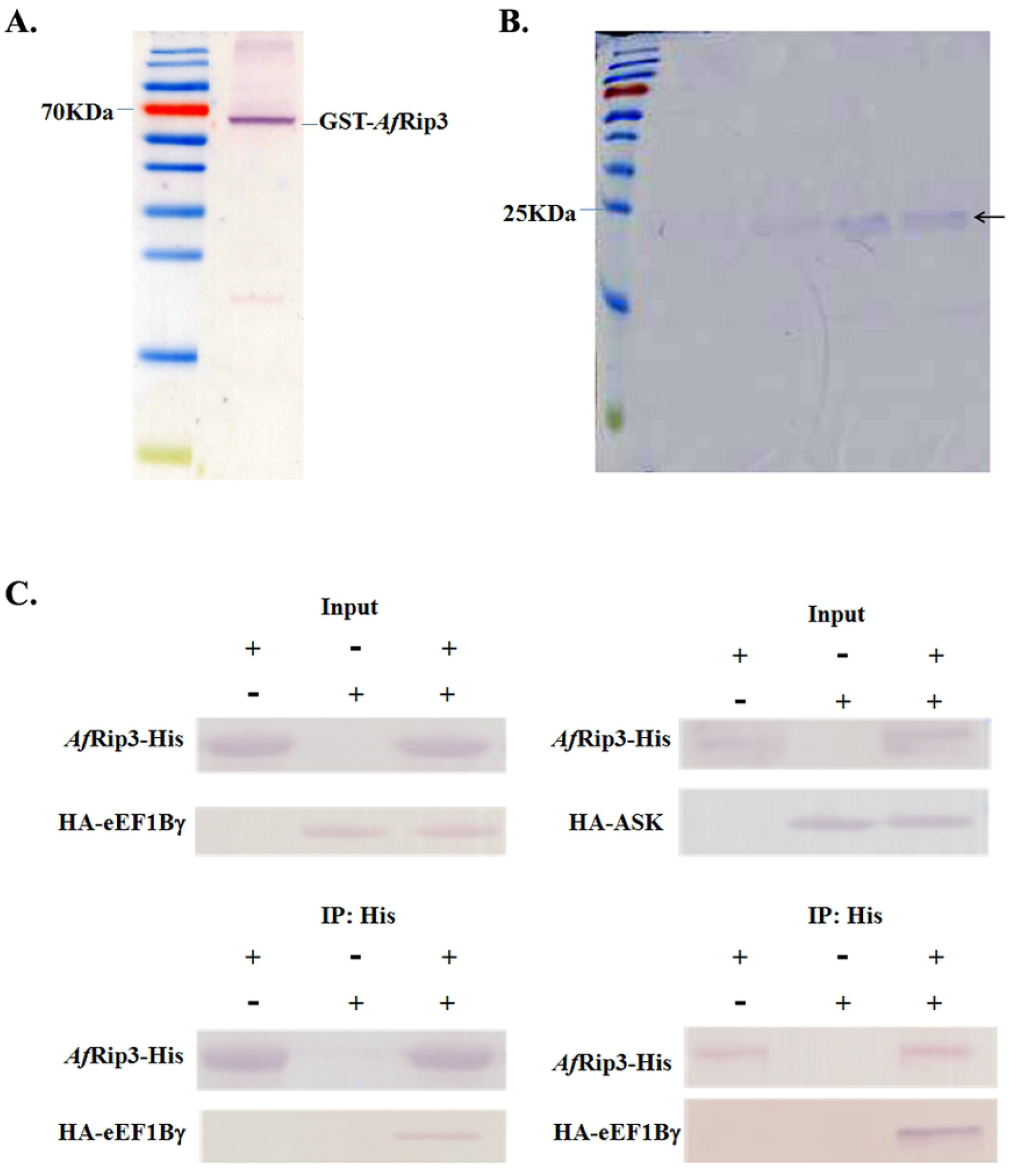
Identification and confirmation of proteins that potentially interact with *Af*Rip3. In A, expression and confirmation of GST-*Af*Rip3 protein in *A. fumigatus* under Materials and Methods; in B, GST-*Af*Rip3 protein was bound to GST™ 4B column with a flow rate of 0.2 mL/min to allow full interaction of GST-*Af*Rip3 protein. Binding proteins were separated on SDS-PAGE and stained with coomassie blue; in C, *Af*Rip3-His was expressed in *E. coli* BL21 (Rossetta) under Materials and Methods. eEF1Bγ or adenylylsulfate kinase (ASK) fused with HA tag was expressed in human embryonic kidney cells 293 under Materials and Methods. Cell lysate was immunoprecipitated with anti-His antibody and subsequently probed with anti-HA antibody.

## Discussion

In mammalian cells, RIP1 and RIP3 are important modulators of apoptosis and necroptosis. When caspase activity is inhibited, RIP1 and RIP3 interact via RHIM to form necrosome, which recruits and activates downstream substrates to trigger necroptosis [8-14]. On the other hand, RIP1-independent cases have been reported recently [15-16]. Over-expression of RIP3 can cause necroptosis of the RIP1^-/-^ and caspase 8^-/-^ murine embryonic fibroblasts (MEFs) [16]. Knock-down of RIP1 did not block cell death when L929 cells were exposed to TNF [49]. When fibroblasts are stimulated by Toll-like receptor 3 (TLR3), elimination of caspase 8 results in RIP3-dependent and RIP1-independent necroptosis [50]. Therefore, it seems that RIP1 is not required for necroptosis under certain conditions.

Previously, in an attempt to identify the necroptotic modulator in *A. fumigatus*, commercially available RIP1 and RIP3 antibodies were used to detect the *A. fumigatus* RIP1 and RIP3 homologs. As a result, a putative protein kinase (AFUA_6G02590), namely *Af*Rip3, was identified by the RIP3 antibody and overexpressed in response to suppression of the GPI anchor synthesis, however, this putative protein kinase only contains a kinase domain, while the homotypic interaction motif (RHIM) that is required for interaction with the RIP1 is absent. It should be pointed out that homologs of *Af*Rip3 were found to be widely distributed in filamentous fungi while no such protein was found in *Saccharomyces cerevisiae*, suggesting its important role in multicellular eukaryotic microbes. On the other hand, no homolog has been detected by the RIP1 antibody [30].

To investigate the role of *Af*Rip3 in *A. fumigatus*, in this study we overexpressed Af*rip3* gene in *A. fumigatus*. As expected the Af*rip3*-overexpressing strain OE-Af*rip3* exhibited an increased cell death even at its log-phase, which confirms that Af*rip3* gene is involved in the cell death of *A. fumigatus*. Analysis of OE-Af*rip3* strain revealed that this increased cell death was triggered via necroptotic pathway, instead of apoptotic or autophagic pathway. Meanwhile a mixture of necroptosis inducers (TNF-α, SM-164 and Z-VAD-FMK) was unable to induce necroptotic process once the Af*rip3* was deleted in *A. fumigatus*. These evidences confirm that *Af*Rip3 plays a central role in necroptotic pathway.

Ca^2+^ is reported to induce autophagic and necroptotic death in fungi and mammals [51]. A connection between Ca^2+^ and necroptosis has been suggested by the observation of the increased intracellular Ca^2+^ concentration upon TNF stimuli [52]. In the meantime, the autophagy pathway initiated by a Ca^2+^-mediated mechanism in some types of cell was also unraveled [53-54]. Also, it is reported that in yeast cell viability was reduced to 80% when incubated with 1 mM or 50 mM Ca^2+^. However, when yeast was incubated with a low concentration of Ca^2+^ such as 0.1 µM Ca^2+^, the cell viability was reduced to 67% [55]. These observations imply that effect of Ca^2+^ on the yeast cell viability is dose-dependent. Our results indicate that Ca^2+^ has a spectacular impact on both autophagic and necroptotic cell death in *A. fumigatus*. We found that the presence of 0.1 mM of Ca^2+^ in culture medium was able to induce expression of the genes not only in necroptotic pathway but also in autophagic pathway. In case of necroptosis, it reasonable to conclude that the increased cytosolic Ca^2+^ is one of factors to activate *Af*Rip3 and then initiates necroptotic cell death in *A. fumigatus.*

In mammalian cells, MLKL has been identified as downstream substrate of RIP3 [56]. However, homolog of MLKL is not found in *A. fumigatus*. To identify the potential downstream substrate of *Af*Rip3, pull-down assay and co-immunoprecipitation were carried out in this study. Based on our results, it is likely that eEF1Bγ and ASK interact with *Af*Rip3. Searching of the DRYGIN (Data Repository of Yeast Genetic INteractions), a database of quantitative genetic interaction network in yeast, with ASK reveals that MET14 is homolog of ASK in *S. cerevisiae*, which is correlated with CSR1, a phosphatidylinositol transfer protein and has a potential role in regulating lipid metabolism under certain conditions. Also MET14 exerts negative genetic interaction on FPR4, a peptidyl-prolyl cis-trans isomerase (PPIase) involved in signal transduction, cell differentiation and apoptosis [57-58]. Presumably, *A. fumigatus* ASK plays a role as MET14 does in yeast, which might be the way that *Af*Rip3 passes the necroptotic signal to its downstream. Somehow, further investigation needs to be carried out.

In summary, in this study we confirmed that *Af*Rip3 was a necroptotic regulator in *A. fumigatus*. Expression of the *Af*Rip3 was induced by Ca^2+^ and interacted with JNK and SMase pathway, which then caused the ROS generation and the rupture of membrane. ASK was identified as potential downstream substrate of *Af*Rip3. Our findings reveal that modulation of necroptotic death of *A. fumigatus* is distinct from that in mammalian cells and may provide a new strategy for development of anti-fungal drug.

## Material and Methods

### Strains and growth conditions

*Aspergillus fumigatus* strain YJ-407 (China General Microbiological Culture Collection Center, CGMCC0386) was maintained on potato glucose (2%) agar slant. *A. fumigatus* strain CEA17 and plasmid pCDA14 are from C. d’Enfert, Institut Pasteur, France. Strain was propagated at 37°C on complete medium (CM), or minimal medium (MM) with 0.5 mM sodium glutamate as a nitrogen source. Uridine and uracil were added at a concentration of 5 mM when required. Mycelia were harvested from strains grown in CM at 37°C with shaking at 200 rpm. At the specified culture time point, mycelia were harvested and washed with distilled water, then frozen in liquid nitrogen and ground. The powder was stored at -80°C for DNA, RNA and protein extraction. Conidia were prepared by growing *A. fumigatus* strains on solid CM with uridine and uracil (CMU) at 37°C for 36 h. The spores were collected, washed twice with 0.01% Tween 20 in PBS and resuspended in PBS, and its concentration was confirmed by haemocytometer counting and viable counting. Vectors and plasmids were propagated in *Escherichia coli* DH5α (Bethesda Research Laboratories).

### Construction of Af*rip3*-overexpressing strain OE-Af*rip3*

The open reading frame of the Af*rip3* was amplified using primer pairs Af*rip3*-up (5’-AGCTTTGTTTAAACATGAATAATGTTCGGCGAAGGCG-3’) and Af*rip3*-down (5’-AGCTTTGTTTAAACGCACTGCCTCCGTCGTCTCA-3’) from *A. fumigatus* cDNA. The PCR products were digested with *Pme*I and ligated into plasmid pVG2.2 (a gift from Leiden University), which contains the *pyrG* as a selective maker. The plasmid obtained (pVG2.2-Af*rip3*) was transformed into *A. fumigatus* CEA17. The OE-Af*rip3* strain was confirmed by PCR amplification. Using Pr-up-1 (5’-ATAGGGCATATTCAACTACCTGGC T-3’) and Pr-down (5’-GTTTA TAGACTCTCAATTCGCGATC-3’) as primers, real-time PCR analysis was carried out to detect a 100-bp fragment of the Af*rip3* gene with 18s rRNA as control.

### Construction of the ΔAf*rip3* mutant

Flanking regions of Af*rip3* gene were amplified from *A. fumigatus* strain YJ-407 genomic DNA. The upstream and downstream flanking regions of Af*rip3* gene were amplified with Up-rip3-5’ (5’-GCGGCCGCGCGCAGAATATGGCCGTGG-3’) and Up-rip3-3’ (5’-GGATCCCCCGGGGACGCCATTGAATCCAGCTC-3’), Down-rip3-5’ (5’-GGATCCCCTGCTTCGCGTTACACCC-3’) and Down-rip3-3’ (5’-ATG CATGCGGCCGCGCACAAGACCGC GACTCGAT-3’), respectively. The amplified fragments were digested with *Bam*HI*/Not*I. The *pyrG*-blaster cassette (8.6 kb) in pCDA14 was obtained by *Hpa*I digestion and cloned into the *Sma*I site between the up- and down-stream non-coding regions of the Af*rip3* to yield pRIP3-pyrG. The resulting plasmid was linearized at a unique *Not*I site and transformed into the CEA17 strain by protoplast transformation [33].

### Caspase activity assay

Proteins were extracted by grounding mycelia in liquid nitrogen, resuspending the powder in ice-cold lysis buffer (50 mM Hepes, pH 7.4, 1 mM DTT, 0.5 mM EDTA, and 0.1% (v/v) Chaps), and centrifugation at 1,500×g for 10 min [29]. The caspase activities of the supernatant against substrates for caspase 8 were determined using a fluorescent assay based on the cleavage of a AMC (7-amino-4-methylcoumarin) dye from the C-terminal of specific peptide substrates (Caspase Fluorescent (AMC) Substrate/Inhibitor QuantiPak™) (BioMol International).

### Necroptosis induction of the ΔAf*rip3* mutant

1×10^6^ spores were inoculated into 1 ml liquid CM with 1 µl TSZ and incubated at 37°C for 4 hours, then stained with 5 µl Hoechst33342 and 5 µl propidium iodide (PI) at 4°C for 20 minutes. The spores were collected and then examined under the fluorescence microscope using a Zeiss Imager A2 (Zeiss, Japan). TSZ, a mixture of TNF-α, SM-164 and Z-VAD-FMK in Necroptosis Inducer Kit (C1058S, Beyotime, China), was used to induce necroptosis. Hoechst and PI were from Apoptosis and Necrosis Assay Kit (C1056, Beyotime, China).

### Real time PCR

Total RNA was isolated with TRIzol reagent (Invitrogen) and 1 μg of RNA samples were reverse-transcribed in a final volume of 20 μL using HiScript II Q RT SuperMix for qPCR (+gDNA wiper) (Vazyme) according to the manufacturer’s instructions. Quantitative RT-PCR was carried out in a CFX960 (Bio-Rad, USA) using the primers (0.5 μM), 1 μL cDNA, 10 µL ChamQ SYBR qPCR Master Mix (Vazyme) in a final volume of 20 μL. Cycle conditions were 95°C for 5 min for the first cycle, followed by 45 cycles of 95°C for 10 s, 60°C for 15 s, and 72°C for 1 s. Quantification of mRNA levels of different genes was performed using the 2^−ΔΔct^ method. Primers used in this study are listed in supplemental S2 Table. Triplicates of samples were analyzed in each assay, and each experiment was repeated at least three times.

### Extraction of extracellular protein

Freshly prepared 2% (w/v) sodium deoxycholate was added into the medium (1/100 in volume), mixed and placed at 4°C for 30 min. Then, 100% trichloroacetic acid (1/10 in volume) was added and reacted at 4°C for 30 min. After centrifugation (15,000 × g for 15 min, 4°C), the extracellular proteins were precipitated, then washed three times with acetone, dried and dissolved.

### Expression and purification of recombinant *Af*Rip3 in *E. coli*

*A. fumigatus* Af*rip3* cDNA was cloned in the pET30a expression vector, in which a stretch of six histidine residues was added at the C-terminal of *Af*Rip3. *E. coli* BL21 (Rossetta) was transformed with the recombinant vector, and protein expression was induced at the log phase of bacterial growth (OD=0.4-0.6) by the addition of IPTG (Sigma-Aldrich) to 0.4 mM for 8 h. The cells were harvested by centrifugation and resuspended in 50 mL 1× binding buffer (0.02 M sodium phosphate, 0.5 M NaCl, 80 mM imidazole, pH 8.0). After sonication, the cell lysate was collected by centrifugation (17500×g for 30 min at 4°C), filtered through a 0.45 mm membrane, and run on a HiTrap chelating HP column (Amersham Pharmarcia Biotech). After washing with 20 column-volumes of binding buffer, the recombinant protein was eluted with a gradient of imidazole (80-500 mM) and dialysed against 50 mM Tris buffer (pH 7.6). The purity of the recombinant protein was judged by SDS-PAGE. The protein concentration was determined by the Bradford assay [59].

### Western blotting

Mycelia were harvested and cellular proteins were extracted with lysis buffer (100 mM Tris-HCl, 0.01% SDS, 1mM DTT, pH7.5). The supernatants were collected after centrifugation (13,000 rpm at 4°C for 10 min) and boiled for 5 min together with 1×loading buffer to perform SDS-PAGE. Subsequently, separated proteins were further transferred onto PVDF membranes (Millipore). The membrane was blocked with 5% fat-free milk in TBST for 2 h at room temperature, and then incubated with appropriate primary antibody at 4°C overnight. Then the membrane was washed three times with TBST buffer and incubated with an AP-conjugated secondary antibody for 1 h at room temperature. After washing three times with TBST buffer, bands were detected with NBT/BCIP reagent.

### Pull-down assay

Overlapping PCR was performed with GST and Af*rip3*. The plasmid pVG2.2-GST-Af*rip3* was constructed in the same way as pVG2.2-Af*rip3*. CEA17 containing pVG2.2-GST-Af*rip3* was cultured in CM for 24 h. Mycelia were harvested and intracellular proteins were extracted. For GST-pulldown experiments, protein extracts were subjected to Glutathione Sepharose 4B. The column was incubated at room temperature for 10 min. The eluate containing the GST-tagged protein was collected and boiled in 1×SDS loading buffer. Protein extracts were separated on a 12% sodium dodecyl sulfate polyacrylamide gel and stained with Coomassie brilliant blue R250. The gel bands were cut from stained gel, destained and subjected to in-gel digestion with trypsin. The digested peptides were desalted with Hypersep C18 SPE cartridge (Thermo Scientific, Bellefonte, PA, USA), precipitated in 100 μL of 0.1% TFA and passed through the conditioned cartridge. The cartridge was washed three times with 1 mL of 0.1% TFA and the peptides were eluted twice with 1 mL of 0.1% TFA in 50% acetonitrile followed by evaporation to dryness in a Speed-Vac. Finally, the sample was resuspended in 0.1% FA (v/v) and analyzed by MALDI-TOF MS using SCIEX TOF/TOF™ 5800 mass spectrometer with laser frequency of 200 Hz at 355 nm. Calibration of the mass spectrometer was performed using standard peptides. 10-20 mg/mL of 2,5-dihydroxybenzoic acid (DHB) was taken as a dot-like matrix. The acceleration voltage was set to 2 kV. For each strain a triplicate was tested.

### Immunoprecipitation

The gene encoding open reading frame of translation elongation factor 1Bγ or adenylylsulfate kinase was amplified with the primer pairs of 5’-CGCGGATCCAT GTCTTTCGGAACAATCTACTCCT/CCGCTCGAGTCAAGCCTTGGGAATCTC ACG-3’ and 5’-CCCAAGCTTATGGCCACAAAATCACCTACCACG-3’/5’-AAA ACTGCAGCTACTCCTTCTTCGGAGGCAAATAC-3’, respectively. The coding regions were subsequently cloned into a eukaryotic expression vector pXJ40-HA via *BamH*I/*Xho*I, *Hind*III/*Pst*I restriction sites to get recombinant vector respectively. Transient expression of HA-eEF1Bγ tag or HA-ASK in HEK293 cells was transfected and cells were collected, lysed with 1× lysis buffer (Cell Signaling Technology), and then lysates were centrifugated and incubated with *Af*Rip3-His protein and anti-His-tag mAb-Magnetic Beads for 4 h. The beads were exposed to a magnetic field, washed three times with IP buffer (50 mM Tris-HCl (pH7.4), 150 mM NaCl and 1% Nonidet P40), boiled in 1×SDS loading buffer for 5 min, and then analyzed by western blotting.

## Acknowledgments

This work was supported by the National Natural Science Foundation of China (31320103901 and 31630016).

## Supporting information captions

**S1 Table. Pull-down proteins identified by Mass Spectrum.** *A. fumigatus* CEA17 was transformed with pVG2.2-GST-Af*rip3* and cultured in CM for 24 h. Mycelia were harvested and intracellular proteins were extracted. Protein extracts were subjected to Glutathione Sepharose 4B. The column was incubated at room temperature for 10 min. The eluate containing the GST-tagged proteins was collected and boiled in 1×SDS loading buffer. Protein extracts were separated on a 12% SDS-PAGE gel and stained with Coomassie brilliant blue R250. The gel bands were cut from stained gel, destained and subjected to in-gel digestion with trypsin. The digested peptides were analyzed by MALDI-TOF MS as described under Materials and methods.

**S2 Table. Primers used in this study.** Quantitative RT-PCR was carried out in a CFX960 (Bio-Rad, USA) using the primers (0.5 μM), 1 μL cDNA, 10 µL ChamQ SYBR qPCR Master Mix (Vazyme) in a final volume of 20 μL. Cycle conditions were 95°C for 5 min for the first cycle, followed by 45 cycles of 95°C for 10 s, 60°C for 15 s, and 72°C for 1 s. Quantification of mRNA levels of different genes was performed using the 2^−ΔΔct^ method. Triplicates of samples were analyzed in each assay, and each experiment was repeated at least three times.

## Reference

1. Yang Z and Klionsky DJ. Mammalian autophagy: core molecular machinery and signaling regulation. Curr Opin Cell Biol. 2010; 22(2): 124–131.

2. Proskuryakov SY, Konoplyannikov AG, Gabai VL. Necrosis: a specific form of programmed cell death? Exp Cell Res. 2003; 283(1): 1–16.

3. Golstein P and Kroemer G. Cell death by necrosis: towards a molecular definition. Trends Biochem Sci. 2007; 32(1): 37–43.

4. Festjens N, Vanden Berghe T, Vandenabeele P. Necrosis, a well-orchestrated form of cell demise: signalling cascades, important mediators and concomitant immune response. Biochim Biophys Acta. 2006; 1757(9-10): 1371–1387.

5. Cho YS, Challa S, Moquin D, Genga R, Ray TD, Guildford M, et al. Phosphorylation-driven assembly of the RIP1-RIP3 complex regulates programmed necrosis and virus-induced inflammation. Cell. 2009; 137(6): 1112–1123.

6. Scaffidi P, Misteli T, Bianchi ME. Release of chromatin protein HMGB1 by necrotic cells triggers inflammation. Nature. 2002; 418(6894): 191–195.

7. Vanden Berghe T, Kalai M, Denecker G, Meeus A, Saelens X, Vandenabeele P. Necrosis is associated with IL-6 production but apoptosis is not. Cellular Signalling. 2006; 18(3): 328–335.

8. Sridharan H and Upton JW. Programmed necrosis in microbial pathogenesis. Trends Microbiol. 2014; 22(4): 199–207.

9. Yu PW, Huang BC, Shen M, Quast J, Chan E, Xu X, et al. Identification of RIP3, a RIP-like kinase that activates apoptosis and NF kappa B. Current Biology. 1999; 9(10): 539–542.

10. Li J, McQuade T, Siemer AB, Napetschnig J, Moriwaki K, Hsiao YS, et al. The RIP1/RIP3 necrosome forms a functional amyloid signaling complex required for programmed necrosis. Cell. 2012; 150(2): 339–350.

11. He SD, Wang L, Miao L, Wang T, Du F, Zhao L, et al. Receptor interacting protein kinase-3 determines cellular necrotic response to TNF-alpha. Cell. 2009; 137(6): 1100–1111.

12. He S, Huang S, Shen Z. Biomarkers for the detection of necroptosis. Cell Mol Life Sci. 2016; 73(11-12):2177–2181.

13. Sun L, Wang H, Wang Z, He S, Chen S, Liao D, et al. Mixed lineage kinase domain-like protein mediates necrosis signaling downstream of RIP3 kinase. Cell. 2012; 148(1-2): 213–227.

14. Wang H, Sun L, Su L, Rizo J, Liu L, Wang LF, et al. Mixed lineage kinase domain-like protein MLKL causes necrotic membrane disruption upon phosphorylation by RIP3. Mol Cell. 2014; 54(1): 133–46.

15. Zhang DW, Shao J, Lin J, Zhang N, Lu BJ, Lin SC, et al. RIP3, an energy metabolism regulator that switches TNF-induced cell death from apoptosis to necrosis. Science. 2009; 325(5938): 332–336.

16. Moujalled DM, Cook WD, Okamoto T, Murphy J, Lawlor KE, Vince JE, et al. TNF can activate RIPK3 and cause programmed necrosis in the absence of RIPK1. Cell Death Disease. 2013; 4:e465.

17. Latgé JP. The pathobiology of *Aspergillus fumigatus*. Trends Microbiol. 2001; 9: 382–389.

18. Latgé, J. P. *Aspergillus fumigatus* and aspergillosis. Clin Microbiol Rev. 1999; 12: 310–350.

19. Zmeili OS and Soubani AO. Pulmonary aspergillosis: a clinical update. Q J Med. 2007; 100: 317–334.

20. Brown GD, Denning DW, Gow NA, Levitz SM, Netea MG, White TC. Hidden killers: human fungal infections. Sci Transl Med. 2012; 4: 165rv113.

21. Walsh TJ, Anaissie EJ, Denning DW, Herbrecht R, Kontoyiannis DP, Marr KA, et al. Treatment of aspergillosis: clinical practice guidelines of the Infectious Diseases Society of America. Clin Infect Dis. 2008; 46: 327–360

22. Anderson JB. Evolution of antifungal-drug resistance: mechanisms and pathogen fitness. Nat Rev Microbiol. 2005; 3: 547–556.

23. Spanakis EK, Aperis G, Mylonakis E. New agents for the treatment of fungal infections: clinical efficacy and gaps in coverage. Clin Infect Dis. 2006; 43: 1060–1068.

24. Denning DW and Bromley MJ. How to bolster the antifungal pipeline. Science. 2015; 347(6229): 1414–1416.

25. Matsuyama S, Nouraini S, Reed JC. Yeast as a tool for apoptosis research. Curr Opin Microbiol. 1999; 2(6): 618–23.

26. Lam E. Controlled cell death, plant survival and development. Nat Rev Mol Cell Biol. 2004; 5(4): 305–315.

27. Golstein P, Aubry L, Levraud JP. Cell-death alternative model organisms: why and which? Nat Rev Mol Cell Biol. 2003; 4(10): 798–807.

28. Engelberg-Kulka H, Amitai S, Kolodkin-Gal I, Hazan R. Bacterial programmed cell death and multicellular behavior in bacteria. Plos Genetics. 2006; 2(10): 1518–1526.

29. Mousavi SAA and Robson GD. Entry into the stationary phase is associated with a rapid loss of viability and an apoptotic-like phenotype in the opportunistic pathogen *Aspergillus fumigatus*. Fung Genet Biol. 2003; 39(3): 221–229.

30. Yan J, Du T, Zhao W, Hartmann T, Lu H, Lü Y, et al. Transcriptome and biochemical analysis reveals that suppression of GPI-anchor synthesis leads to autophagy and possible necroptosis in *Aspergillus fumigatus*. PlosOne. 2013; 8(3).

31. Meyer V, Wanka F, van Gent J, Arentshorst M, van den Hondel C A, Ram AF. Fungal gene expression on demand: an inducible, tunable, and metabolism-independent expression system for *Aspergillus niger*. Appl Environ Microbiol. 2011; 77(9): 2975–2983.

32. Vogt K, Bhabhra R, Rhodes JC, Askew DS. Doxycycline-regulated gene expression in the opportunistic fungal pathogen *Aspergillus fumigatus*. BMC Microbiol. 2005; 5: 1, DOI: 10.1186/1471-2180-5-1.

33. Yelton MM, Hamer JE, Timberlake WE. Transformation of *Aspergillus nidulans* by using a trpC plasmid. Proc Natl Acad Sci U S A. 1984; 81(5): 1470–1474.

34. Brana C, Benham C, Sundstrom L. A method for characterising cell death in vitro by combining propidium iodide staining with immunohistochemistry. Brain Research Protocols. 2002; 10(2): 109–114.

35. Hailer NP, Vogt C, Korf HW, Dehghani F. Interleukin-1 beta exacerbates and interleukin-1 receptor antagonist attenuates neuronal injury and microglial activation after excitotoxic damage in organotypic hippocampal slice cultures. Eur J Neurosci. 2005; 21(9): 2347–2360.

36. Miller LDP, Mahanty NK, Connor JA, Landis DM. Spontaneous pyramidal cell-death in organotypic slice cultures from rat hippocampus is prevented by glutamate-receptor antagonists. Neurosci. 1994; 63(2): 471–487.

37. Danial NN and Korsmeyer SJ. Cell death: Critical control points. Cell. 2004; 116(2): 205–219.

38. Gardai SJ, Bratton DL, Ogden CA, Henson PM. Recognition ligands on apoptotic cells: a perspective. J Leukoc Biol. 2006; 79(5): 896–903.

39. Eskelinen EL. New insights into the mechanisms of macroautophagy in mammalian cells. Int Rev Cell Mol Biol. 2008; 266: 207–247.

40. Xie X, Zhao Y, Ma CY, Xu XM, Zhang YQ, Wang CG, et al. Dimethyl fumarate induces necroptosis in colon cancer cells through GSH depletion/ROS increase/MAPKs activation pathway. British J Pharmacol. 2015; 172(15): 3929–3943.

41. Tanida I. Autophagosome formation and molecular mechanism of autophagy. Antioxid Redox Signal. 2011; 14(11): 2201–2214.

42. Ventura JJ, Cogswell P, Flavell RA, Baldwin AS Jr, Davis RJ. JNK potentiates TNF-stimulated necrosis by increasing the production of cytotoxic reactive oxygen species. Genes Dev. 2004; 18(23): 2905–2915.

43. Xu Y, Huang S, Liu ZG, Han J. Poly(ADP-ribose) polymerase-1 signaling to mitochondria in necrotic cell death requires RIP1/TRAF2-mediated JNK1 activation. J Biol Chem. 2006; 281(13): 8788–8795.

44. Antosiewicz J, Ziolkowski W, Kaczor JJ, Herman-Antosiewicz A. Tumor necrosis factor-alpha-induced reactive oxygen species formation is mediated by JNK1-dependent ferritin degradation and elevation of labile iron pool. Free Radic Biol Med. 2007; 43(2): 265–270.

45. Won JS and Singh I. Sphingolipid signaling and redox regulation. Free Radic Biol Med. 2006; 40(11): 1875–1888.

46. Mao C and Obeid LM. Ceramidases: regulators of cellular responses mediated by ceramide, sphingosine, and sphingosine-1-phosphate. Biochim Biophys Acta. 2008; 1781(9): 424–34.

47. Orrenius S, Gogvadze V, Zhivotovsky B. Mitochondrial oxidative stress: implications for cell death. Annu Rev Pharmacol Toxicol. 2007; 47: 143–183.

48. He SD, Liang Y, Shao F, Wang X. Toll-like receptors activate programmed necrosis in macrophages through a receptor-interacting kinase-3-mediated pathway. Proc Natl Acad Sci U S A. 2011; 108(50): 20054–20059.

49. Vanlangenakker N, Bertrand MJ, Bogaert P, Vandenabeele P, Vanden Berghe T. TNF-induced necroptosis in L929 cells is tightly regulated by multiple TNFR1 complex I and II members. Cell Death Disease. 2011; 2 :e230, doi: 10.1038/cddis.2011.111.

50. Kaiser WJ, Sridharan H, Huang C, Mandal P, Upton JW, Gough PJ, et al. Toll-like receptor 3-mediated necrosis via TRIF, RIP3, and MLKL. J Biol Chem. 2013; 288(43): 31268–31279.

51. Khan MJ, Rizwan AM, Waldeck-Weiermair M, Karsten F, Groschner L, Riederer, M, et al. Inhibition of autophagy rescues palmitic acid-induced necroptosis of endothelial cells. J Biol Chem. 2012; 287(25): 21110–21120.

52. Ono K, Kim SO, Han JH. Susceptibility of lysosomes to rupture is a determinant for plasma membrane disruption in tumor necrosis factor alpha-induced cell death. Mol Cell Biol. 2003; 23(2): 665–676.

53. Crawford SE and Estes MK. Viroporin-mediated calcium-activated autophagy. Autophagy. 2013; 9(5): 797–8.

54. Medina DL and Ballabio A. Lysosomal calcium regulates autophagy. Autophagy. 2015; 11(6): 970–971.

55. Halachmi D and Eilam Y. Calcium homeostasis in yeast cells exposed to high concentration of calcium: Roles of vacuolar H-ATPase and cellular ATP. FEBS Lett. 1993; 316(1): 73–78.

56. Murphy JM, Czabotar PE, Hildebrand JM, Lucet IS, Zhang JG, Alvarez-Diaz S, et al. The pseudokinase MLKL mediates necroptosis via a molecular switch mechanism. Immunity. 2013; 39(3): 443–453.

57. Schiene-Fischer C. Multidomain peptidyl prolyl cis/trans isomerases. Biochim Biophys Acta. 2015; 1850(10): 2005–2016.

58. Nakagawa T, Shimizu S, Watanabe T, Yamaguchi O, Otsu K, Yamagata H, et al. Cyclophilin D-dependent mitochondrial permeability transition regulates some necrotic but not apoptotic cell death. Nature. 2005; 434(7033): 652–658.

59. Bradford MM. A rapid and sensitive method for the quantitation of microgram quantities of protein utilizing the principle of protein-dye binding. Anal Biochem. 1976; 72: 248–254.

